# Control of Genetic Parasites Mediated Through Nucleoporin Evolution

**DOI:** 10.1101/215913

**Authors:** Paul A. Rowley, Kurt Patterson, Suzanne B. Sandmeyer, Sara L. Sawyer

## Abstract

Yeasts serve as long-term hosts to several types of genetic parasites. Few studies have addressed the evolutionary trajectory of yeast genes that control the stable co-existence of these parasites with their host cell. In *Saccharomyces* yeasts, the retrovirus-like Ty retrotransposons must access the nucleus. We show that several genes encoding components of the yeast nuclear pore complex have experienced natural selection for substitutions that change the encoded protein sequence. By replacing these *S. cerevisiae* genes with orthologs from other *Saccharomyces* species, we discovered that natural sequence changes have affected the control of Ty retrotransposons. Specifically, changing the genetic sequence of *NUP84* or *NUP82* to what is found in other *Saccharomyces* species alters the retrotransposition of *S. cerevisiae* Ty1 and Ty3, respectively. Importantly, all tested housekeeping functions of *NUP84* and *NUP82* remained equivalent across species. The nuclear pore complex is the gatekeeper of the nucleus. It appears that nucleoporins are adapting to modulate the control of genetic parasites which access the nucleus, which is achieved despite the strict constraints imposed by host nuclear pore complex function.

---

The presence of Ty retrotransposons (Tys) in all species of *Saccharomyces* yeasts suggest that they have likely been coevolving together for about 20 million years [1,2]. Because Tys are strictly intracellular parasites, both the host (yeast) and Tys are aligned in benefitting from a controlled, sustained relationship that does not place the host at an evolutionary disadvantage [3]. This might even be thought of as a symbiotic relationship because, unlike most pathogenic viruses of higher eukaryotes, Tys are a force for genetic plasticity, driving adaptive changes within the yeast genome in response to changes in environmental conditions [4]. For this reason, it is thought that both Tys and the host genome have evolved mechanisms to attenuate unchecked Ty replication that would place an excessive burden on the host cell [3,5–10]. Thus, yeasts have likely experienced selection to control genetic parasites [11,12]. In turn, Tys may counter-adapt to host control strategies, or may adapt to modulate their own pathogenicity. Regardless of whether a Ty is thought of as a symbiont, or a “tamed” parasite, one can imagine that the host-parasite relationship must be finely honed within each yeast species, with different evolutionary strategies emerging over evolutionary time (in both yeast and Ty) to control Ty replication.

There are many examples of genetic parasites, including viruses and transposable elements, that must access the nucleus of a host cell in order to replicate. Thus, the nuclear envelope and nuclear gating represents a major barrier to these parasites in their eukaryotic hosts [13–15]. The movement of large macromolecules between the cytoplasm and the nucleus occurs though the nuclear pore complex. The nuclear pore complex is composed of multiple copies of approximately 30 different proteins, referred to as nucleoporins, and is conserved between yeast and higher eukaryotic species, including humans [16–22]. Transport receptors, called karyopherins, facilitate the transport of cellular cargo through the nuclear pore [20,23]. Genetic parasites interact with a wide variety of nucleoporins and karyopherins to facilitate the nucleocytoplasmic transport of their proteins and complexes, or to re-localize useful or harmful host proteins [24–33].

*Saccharomyces* yeasts are eukaryotes chronically infected with DNA plasmids, single-stranded RNA viruses, double stranded RNA viruses, and retrotransposons [34,35]. Of these virus and virus-like elements, only Tys transit through the nuclear pore complex. There are five families of Tys in *S. cerevisiae,* Ty1 to Ty5, and all have an analogous lifecycle to mammalian retroviruses [36–38]. Tys have intracellular lifecycles (Figure 1), but can be transmitted to new hosts via mating [39]. The Ty lifecycle involves the shuttling of Ty RNAs (with associated host and viral proteins) in and out of the nucleus every replication cycle. Ty3 virus-like particles and proteins have been observed to cluster at the nuclear envelope and the cytoplasmic face of the nuclear pore complex [25,40,41]. Multiple Ty3 proteins (Gag3, p27 and CA) interact directly with nucleoporins, and the Ty1 and Ty3 integrase (IN) proteins contain nuclear localization signals [25,42–45]. Together, these factors presumably direct the nuclear ingress of Ty cDNA and associated proteins. After nuclear entry, integrase catalyzes the insertion of Ty cDNA into the host genome [46,47]. Tys must also exit the nucleus. Ty1 RNAs, after transcription in the nucleus, are thought to be stabilized and chaperoned from the nucleus by the Gag protein [40].

**Figure 1.**
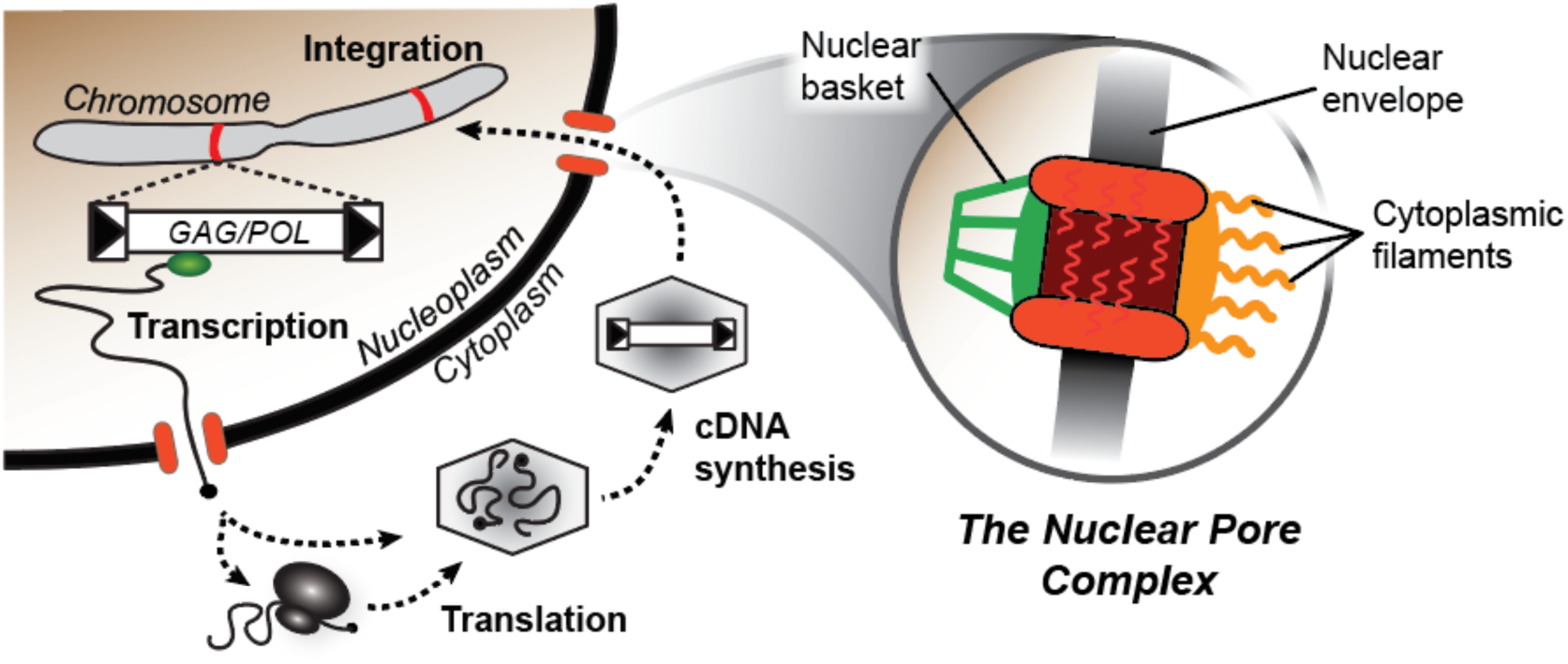
The nuclear pore complex is important for Ty retrotransposition. *Left.* A generic schematic of the lifecycle of a Ty. Chromosomal copies of Ty, found in the yeast genome, produce full-length RNA transcripts that are exported from the nucleus. These transcripts are then translated and packaged within virus-like particles within the cytoplasm. Packaged RNAs are reverse transcribed into cDNA that is transported into the nucleus via the nuclear pore complex. The Ty integrase mediates insertion of the cDNA into the host genome at a new location (red stripes on the chromosome). *Right.* Simplified representation of the nuclear pore complex embedded in the nuclear envelope and sliced along its vertical axis. Filaments rich in phenylalanine and glycine (FG) radiate into the nucleoplasm, cytoplasm, and within the nuclear pore itself.

Because the lifecycle of Tys involves trafficking in and out of the nucleus, we investigated the hypothesis that nucleoporins might experience evolutionary pressure to control Ty nucleocytoplasmic transport. While evolution of host immune strategies is common [48–50], evolved resistances have not been extensively documented in large, essential cellular assemblages, such as the nuclear pore complex. At least six published high-throughput gene knockout screens have been conducted in order to identify genes important for the replication of Ty1 (four studies [51–54]) or Ty3 (two studies [55,56]). Among these studies, nine nucleoporins (Figure 2A) and four karyopherins (Figure S1) were identified as important for Ty replication. Several genes were identified in multiple screens, as represented in the Venn diagrams shown in Figures 2A and S1. Interestingly, the knockout of some nuclear pore-related genes has been noted to reduce Ty retrotransposition, while the knockout of others increase it [57]. One possible interpretation of this confusing pattern is that there is a highly evolved relationship between yeasts and Tys. In some cases, Tys are successfully exploiting a nuclear pore protein for import/export. Knockout of such genes would reduce Ty retrotransposition. In other cases the host may have evolved to reduce Ty replication, for instance by evolving a nuclear pore protein that binds but does not transit Ty componentry, or that binds Ty componentry and mis-localizes it. Deleting these genes would increase Ty replication. There are likely to be additional nuclear pore complex-related genes, beyond those shown in Figures 2A and S1, that are involved in Ty replication. This is because genes essential to yeast viability are likely underrepresented in screens, given that gene knockouts of these are inviable.

**Figure 2.**
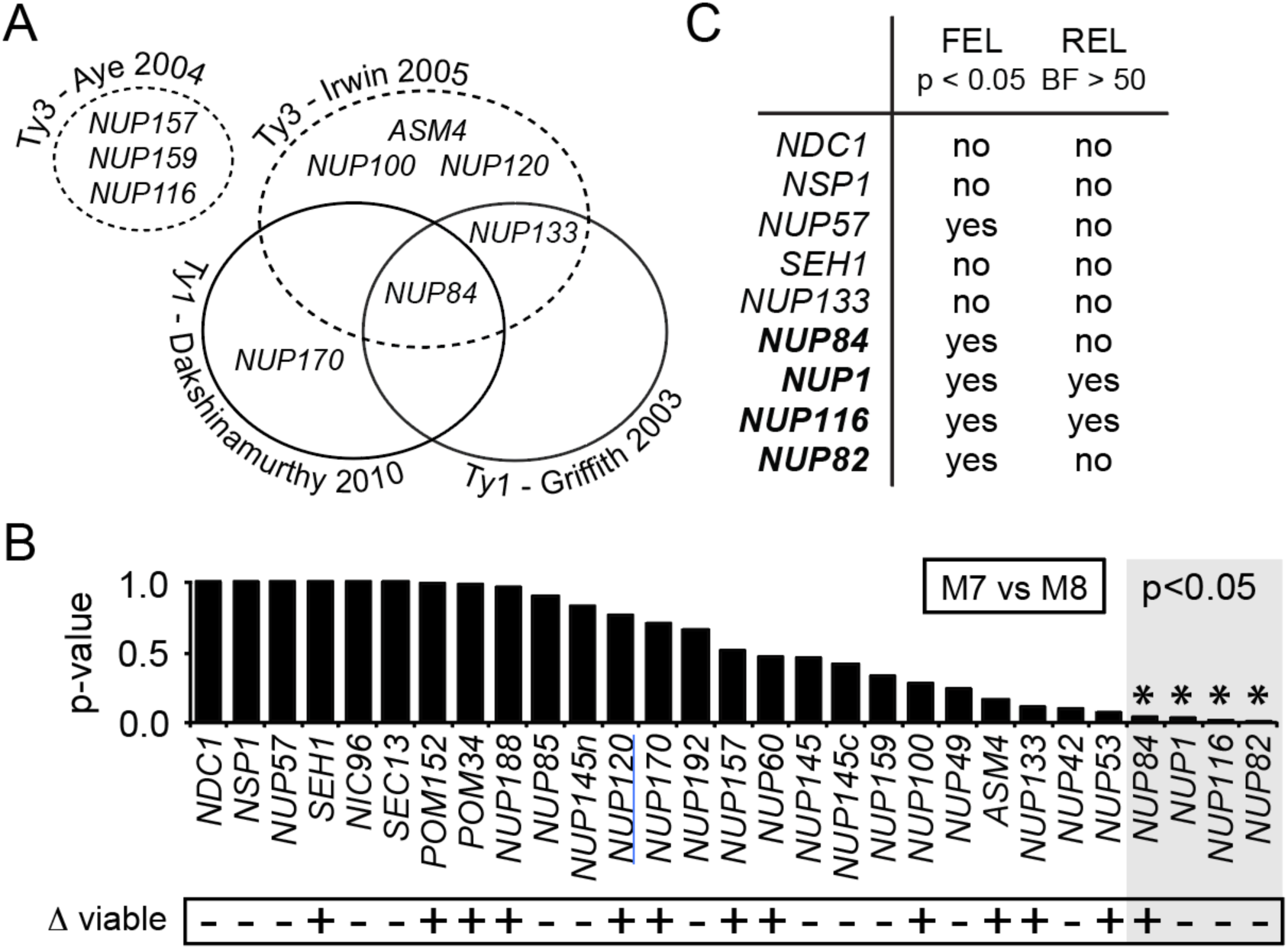
Several components of the nuclear pore complex are evolving rapidly in Saccharomyces yeasts. (A) A Venn diagram shows the results of published high-throughput genetic screens for host factors affecting Ty retrotransposition [51,52,55,56]. Only nucleoporin genes found in these screens are summarized, where disruption of the indicated gene altered Ty3 (dashed lines) or Ty1 (solid lines) retrotransposition. (B) Results from PAML analysis surveying nucleoporin genes for codons with elevated evolutionary rate (dN/dS ≥1). Here, alignments were fit to a codon model of purifying selection (M7) and a codon model allowing for codons with an elevated evolutionary rate (M8). M7 was rejected in favor of M8 for four nucleoporins (p<0.05): *NUP84, NUP1, NUP116* and *NUP82.* Along the bottom is summarized whether yeast with a deletion of each of these genes is viable, taken from the *Saccharomyces* genome database. (C) Extended evolutionary analysis of selected nucleoporins using two common tests for positive selection (FEL and REL) [67]. “Yes” indicates that codons with dN/dS>1 were detected in this gene by the indicated test, with a p-value (p) < 0.05, or Bayes factor (BF) > 50, as indicated across the top.

To further explore the idea of evolved control of Tys, we looked at the evolutionary history of all known *Saccharomyces* nucleoporin genes, and found that 26 of 30 nucleoporins have changed very little during *Saccharomyces* speciation and are evolving under purifying selection. However, four nucleoporins are evolving rapidly in a manner consistent with positive selection (*NUP1, NUP82, NUP84,* and *NUP116).* We wished to explore how this high level of protein-level sequence divergence between species would affect Ty control. For *NUP82* and *NUP84,* we engineered *S. cerevisiae* strains to express orthologs from other yeast species and then assayed the replicative success of different families of Tys within these otherwise isogenic yeast strains. We found that species-specific evolutionary differences in these nucleoporins affected the replication of either Ty1, Ty3, or both Ty familie*S. NUP84*appears to have experienced selection primarily to limit Ty1, while *NUP82* has experienced selection primarily to limit Ty3. Moreover, Nup82p and Nup84p are integral to the nuclear pore complex structure that are essential for its proper functioning [58,59]. We find that these adaptive changes in *NUP82* and *NUP84* affect Ty replication, yet have accumulated under the constraints of strict conservation of nucleoporin host functions throughout *Saccharomyces* speciation.

## RESULTS

### *NUP82* and *NUP84* have accumulated elevated levels of non-synonymous mutations

We first set out to determine which nuclear pore complex-related genes might be important in the evolved control of Tys. Obviously, genes that have remained unchanged over the speciation of *Saccharomyces* yeast would be unlikely to fall into this class. Instead, as a screening tool we sought genes that have diverged significantly in sequence from one yeast species to the next. We are particularly interested in genes with evidence for natural selection underlying these sequence changes, rather than genes that have diverged in sequence simply by the forces of random genetic drift. Natural selection can be detected in genes as follows. Typically, selection operates on non-synonymous substitutions (changing the encoded amino acid) more significantly than on nonsynonymous mutations (silent, not changing the encoded amino acids). Gene regions that have experienced repeated rounds of natural selection in favor of protein-altering mutation therefore exhibit a characteristic inflation of the rate of non-synonymous (dN) DNA substitutions compared to synonymous (dS) substitutions (denoted by dN/dS > 1) [60]. Because non-synonymous mutations occur more often than synonymous mutations by random chance, computational models have been developed that use statistical frameworks to account for these unequal substitution rates [61–63]. The mode of evolution that we are seeking (dN/dS > 1) is considered to be somewhat rare in eukaryotic genes. Instead, most genes experience purifying selection (dN/dS < 1), where protein sequence is conserved over evolutionary time due to the important and complex roles that most proteins play in cellular homeostasis.

We examined the evolution of 29 yeast genes encoding nucleoporins and 22 genes encoding karyopherins for evidence of codons with dN/dS > 1. For each gene, we gathered nucleotide sequences from six divergent *Saccharomyces* species (*S. cerevisiae, S. paradoxus, S. mikatae, S. kudriavzevii, S. arboricolus* and *S. bayanus)* [64–66]. Next, we constructed DNA alignments of the various genes and fit these to two different models of codon evolution using the Phylogenetic Analysis by Maximum Likelihood (PAML) package [63]. Evolutionary model M7 was used as our null model and assumes that all codons within a gene are evolving under purifying selection (dN/dS > 1 not allowed), whereas model M8 allows for some codons to exhibit an elevated evolutionary rate (dN/dS ≥ 1). Model M7 was rejected in favor of M8 (p<0.05) for four nucleoporin genes: *NUP84, NUP1, NUP116* and *NUP82* (Figure 2B). The null model was not rejected for any karyopherins (Figure S1). Interestingly, one of these nucleoporin genes, *NUP84,* is also the only nuclear pore-related gene found in three different knockout screens as important for Ty retrotransposition (Figure 2A). *NUP133,* which was the only other nucleoporin found in more than one of the genetic screens (Figure 2A), is important for both Ty1 and Ty3 retrotransposition, but did not pass the threshold of significance (p=0.11; Figure 2B), and so was not investigated further. The remaining three nucleoporin genes under positive selection *(NUP1, NUP116* and *NUP82)* are essential genes within *S. cerevisiae* (Figure 2B, bottom), and of these, only *NUP116* has been directly shown to be involved with Ty replication [25].

We next evaluated *NUP84, NUP1, NUP116,* and *NUP82* with additional tests for positive selection, FEL and REL [67]. We found that all four nucleoporin genes showed evidence of positive selection using at least one of these additional tests (Figure 2C). Furthermore, three of these genes *(NUP1, NUP82,* and *NUP116)*were previously identified as evolving rapidly in a whole genome evolutionary study of five *Saccharomyces* yeast species performed by Scannell *et al.* [66]. In contrast, *NUP133* and four other nucleoporins with the least support for rejection of the M7 null model *(NDC1, NSP1, NUP57* and *SEH1*; Figure 2B), passed zero or only one of these tests (Figure 2C). We next turned to functionally testing the biological relevance of the observed evolutionary signatures identified within nuclear pore complex-related genes.

### *A novel GFP* reporter of Ty retrotransposition

We first built a quantitative, GFP-based assay system for Ty retrotransposition, which is a variation of a previous assay used in this field [68]. In this system, a plasmid-mounted Ty1 genome from *Saccharomyces cerevisiae* was encoded on the Watson (sense) strand, and was engineered to contain an internal *GFP*gene on the Crick (anti-sense relative to the transcript) strand of the DNA (Figure 3A). To prevent its expression prior to retrotransposition, the *GFP*gene was engineered to contain an antisense intron (on the Watson strand). Thus, only after the full-length Ty1*-GFP* transcript has been spliced, reverse transcribed, and integrated into the *S. cerevisiae* genome can the *GFP* gene be expressed. This is further regulated by the inducible copper-sensitive *CUP1* promoter (Figure 3A). Experiments were performed with two different introns within the *GFP* gene in order to determine which was more efficiently spliced from the transcript produced. The more efficient splicing occurred using the *S. cerevisiae ACT1* intron (ACT1i) (Figure 3B). GFP-positive cells were only detected by flow cytometry after galactose was added to the media to initiate Ty1 transcription, and after subsequent addition of CuSO_4_ to the media to induce expression of the *GFP* reporter (Figure 3C). We tested our Ty1 retrotransposition reporter in isogenic strains deleted for five genes known to be important for efficient Ty1 retrotransposition: BY4741 *xrn1Δ, nup84Δ, nup133Δ, bud22Δ,* and *xrs2D* [51,52]. Indeed, we see a significant decrease in Ty1 retrotransposition in each deletion strain compared to the wild-type BY4741 background (Figure 3D). As a control, we show that a strain deleted for *NUP100,* which is important for Ty3 retrotransposition [55], but not known to be important for Ty1, supports a level of retrotransposition that is not significantly different from that of a wild-type strain (Figure 3D).

**Figure 3.**
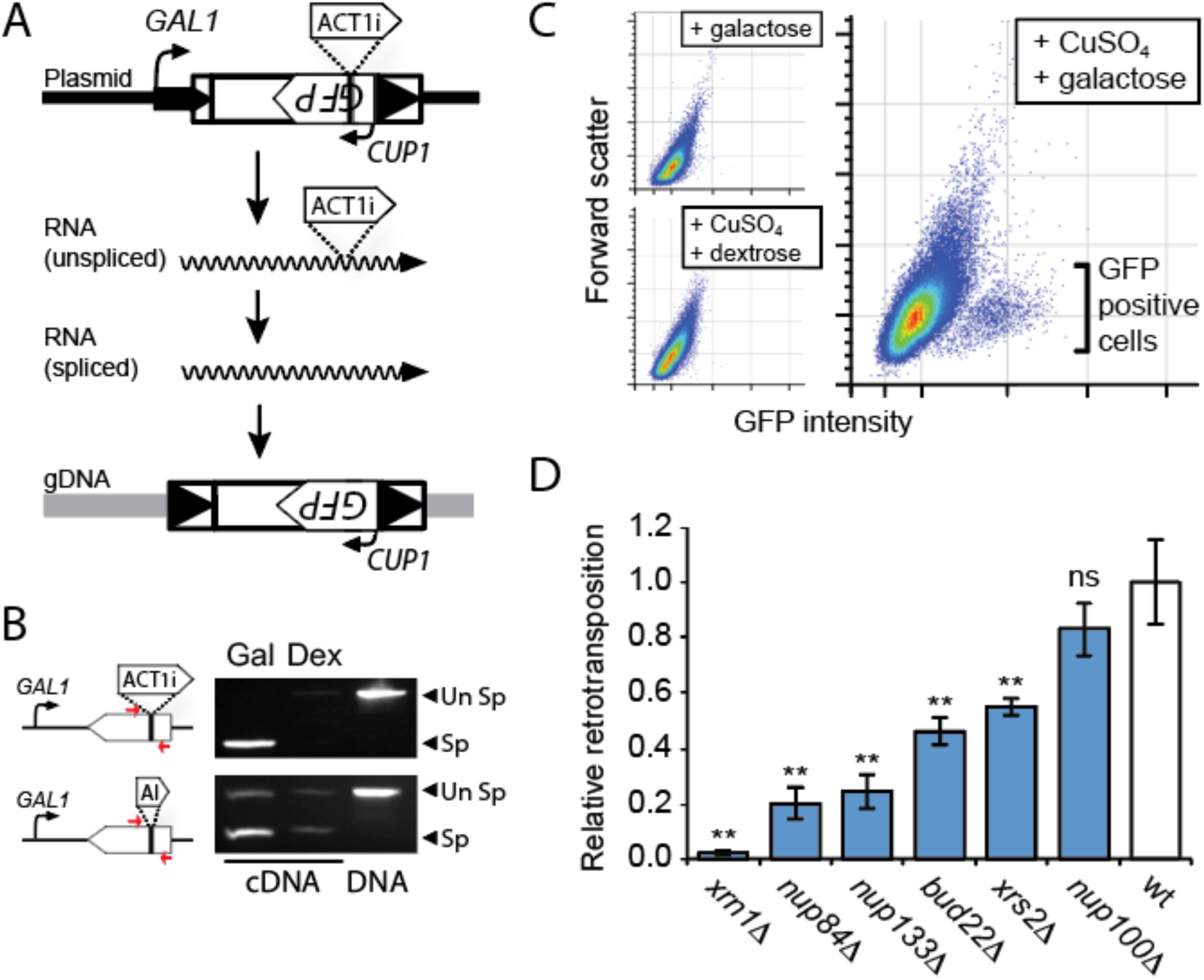
A novel GFP-based reporter of Ty1 retrotransposition. *(A)* An overview of the GFP-tagged Ty1 plasmid. Ty1 transcription is induced by activation of the *GAL1* promoter, and produces a long Ty1 transcript that includes an internal *GFP*gene, and an *ACT1* intron (ACT1i) that is internal to *GFP.* The spliced transcript has ACTi removed, and then provides a template for Ty1 protein production and reverse transcription. At the completion of the Ty1 lifecycle, Ty1 cDNA is imported into the nucleus and integrated into the *S. cerevisiae* genome. The *GFP* gene is then induced from the *CUP1* promoter by CuSO_4_ to report successful integration events. (B) RT-PCR was used to assess splicing of RNA with ACT1i versus an artificial intron (AI) within the *GFP* gene (primer positions marked by red arrows). Spliced RNA transcripts (Sp) were mainly detected upon induction of the transcription by the *GAL1* promoter using galactose (Gal). Growth on dextrose (Dex) inhibits the *GAL1* promoter and the production of RNA transcripts. Plasmid DNA was used as positive control to allow the PCR amplification across intron-containing *GFP.* “Un Sp” is unspliced RNA. (C) Flow cytometry analysis shows that *GFP* is only expressed under conditions of galactose induction of Ty1 expression followed by CuSO_4_ induction of *GFP.* (D) The effect of six different gene deletions on Ty1 retrotransposition, relative to wild-type *S. cerevisiae.* The relative retrotransposition was measured as a percent of GFP positive cells after induction of the Ty-GFP reporter, and was repeated independently, three times (error bars: standard error, n>3; **Tukey-Kramer method, p<0.05). All values are normalized to wildtype.

### *NUP84* evolution modulates Ty1 retrotransposition within *S. cerevisiae*

*NUP84* is under positive selection and disruption of the gene affects both Ty1 and Ty3 replication (Figure 1). We wished to test whether the evolution of *NUP84* over yeast speciation has altered interactions with Tys. To test this, we replaced *NUP84* within the *S. cerevisiae* genome *(NUP84^Scer^)* with *NUP84* from diverse *Saccharomyces* species (*S. mikatae, S. kudriavzevii*and *S. bayanus)* using the method outlined in Figure 4A. These sequences encode Nup84p that are between 91% (S. paradoxus) and 84% (*S. bayanus)* identical to the *S. cerevisiae* protein. As an isogenic control, we re-complemented the *nup84D* strain with *S. cerevisiae NUP84,* as was done for the other *Saccharomyces*orthologs. Chromosomal complementation of *S. cerevisiae nup84D* with each heterospecific (other species) *NUP84* allele resulted in the restoration of normal growth and cellular morphology (Figure 4B and 4D), normal nuclear import (Figure 4C and 4D), and normal transcription from the promoters used in our Ty1 GFP-based reporter (Figure 4E).

**Figure 4.**
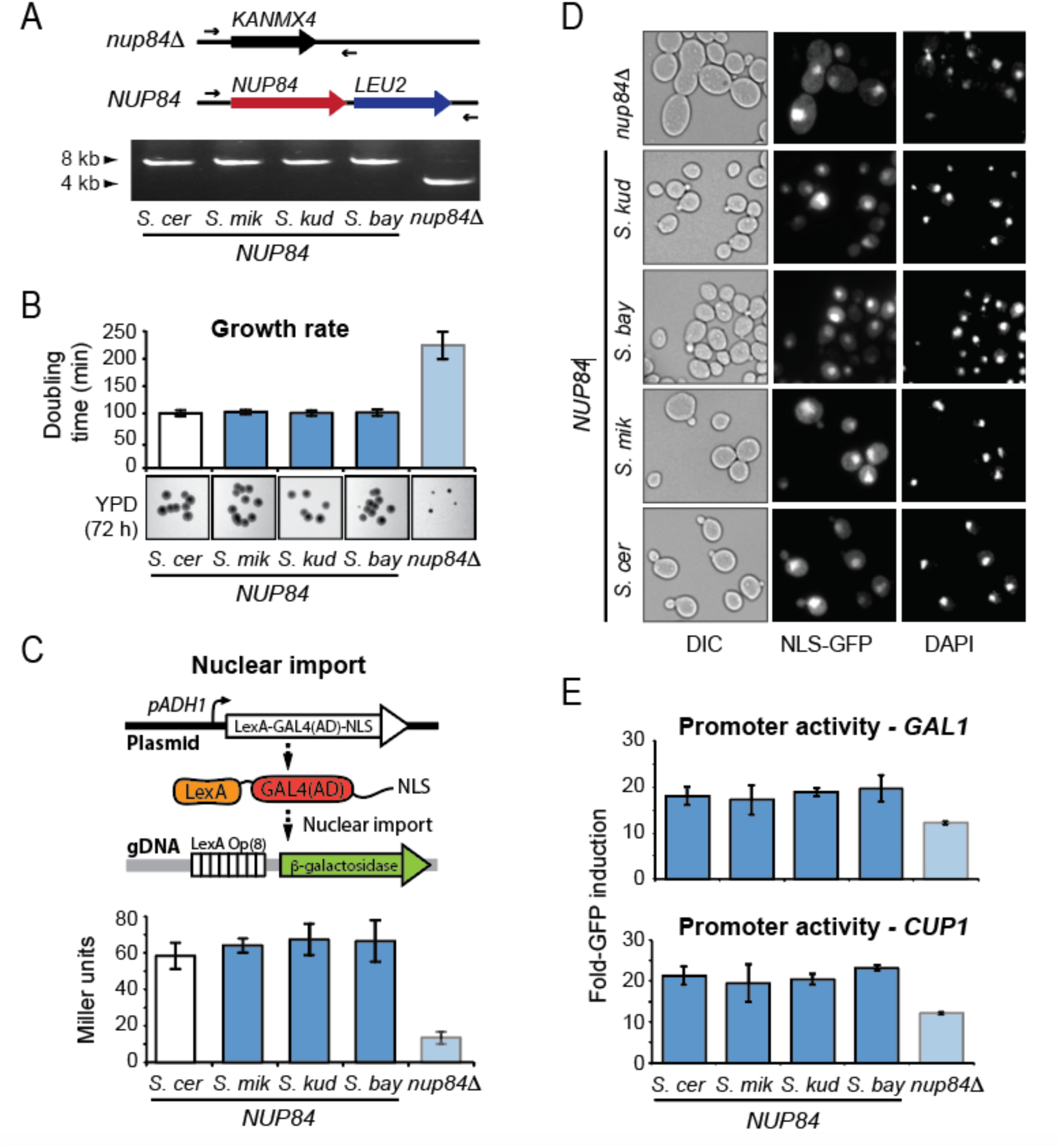
The housekeeping functions of NUP84 areconserved across divergent Saccharomycesspecies. (A) *Top.* A schematic representation of the *NUP84*locus within *S. cerevisiae* engineered to either lack *NUP84 (nup84D)* or to express heterospecific *NUP84*from *S. cerevisiae (S. cer), S. mikatae (S. mik), S. kudriavzevii* (*S. kud)* or *S. bayanus (S. bay)* along with the *LEU2* selectable marker. Successful genome engineering was confirmed by the PCR amplification (primers marked as arrows) across the *NUP84*locus to detect the replacement of *KANMX4* with *NUP84-LEU2.* (B) The doubling time of NUP84-complemented strains in liquid YPD medium compared to *nup84D,* and colony growth and morphology after 72 h of growth on solid YPD medium. (C) General nuclear import function was assessed in the presence of heterospecific Nup84p or absence of Nup84p using a LexA-Gal4(AD) reporter protein with a SV40 nuclear localization signal (NLS) [69]. The LexA DNA binding domain and Gal4 activation domain (AD) initiate transcription of the β-galactosidase gene upon successful nuclear import. (D) Nuclear transport was also assessed by the steady-state localization of a GFP reporter protein containing a NLS from *PHO4* [70] and its cellular accumulation relative to a DAPI-stained nucleus within *NUP84* complemented *S. cerevisiae.* (E) The effect of *NUP84* complementation or deletion on the ability of *S. cerevisiae*to express *GFP* from each of the promoters used in the Ty1 GFP-based reporter *(GAL1*(top) or *CUP1* (bottom) promoters), using mean fluorescent intensity (MFI) detected by flow cytometry (error bars: standard error, n>3).

The null strain, and each of the four strains expressing wildtype of heterospecific *NUP84,* were transformed with the Ty1 *GFP* reporter described above. Relative to *nup84D,* cells complemented with *NUP84*^*S.cer*^ increased Ty1 retrotransposition approximately 5-fold (Figure 5A). There were highly significant differences in the levels of retrotransposition among strains encoding heterospecific *NUP84* (one-way ANOVA, p=8.2 × 10^−8^), and levels of Ty1 retrotransposition were significantly different in strains containing *NUP84*^*S.mik*^, *NUP84*^*S.kud*^, and *NUP84*^*Sbay*^ when compared to *NUP84*^*S.cer*^(Tukey-Kramer method, p<0.05) (Figure 5A). We found that replacement of *NUP84*^*S.cer*^ with *NUP84*^*S.kud*^ increased Ty1 retrotransposition by 32%, whereas *NUP84*^*S.mik*^ and *NUP84*^*S.bay*^ both significantly decreased retrotransposition by 21% and 35%, respectively. To verify the observed differences in control of Ty1 retrotransposition, we used Southern blotting to detect Ty1 integrations in the 5’ UTR of the *SUF16* locus, as previously described [71]. We used our *GFP* reporter assay to initiate Ty1 retrotransposition, with Ty1 genomic integrations only detected after induction by galactose (Figure 5B). Similar to our *GFP* reporter assay, fewer integrations were detected within strains encoding *NUP84*^*S mik*^ and *NUP84*^*S.bay*^ compared to *NUP84*^*S.cer*^. *NUP84*^*S.cer*^ and *NUP84*^*S.kud*^ had comparable levels of genomic integrations (Figure 5B). It is important to remember that these assays are conducted over a short period, yet Ty expand over time [5,6]. Therefore, even small differences in these assays would be expected to have significant fitness impacts on the host and on Ty.

**Figure 5.**
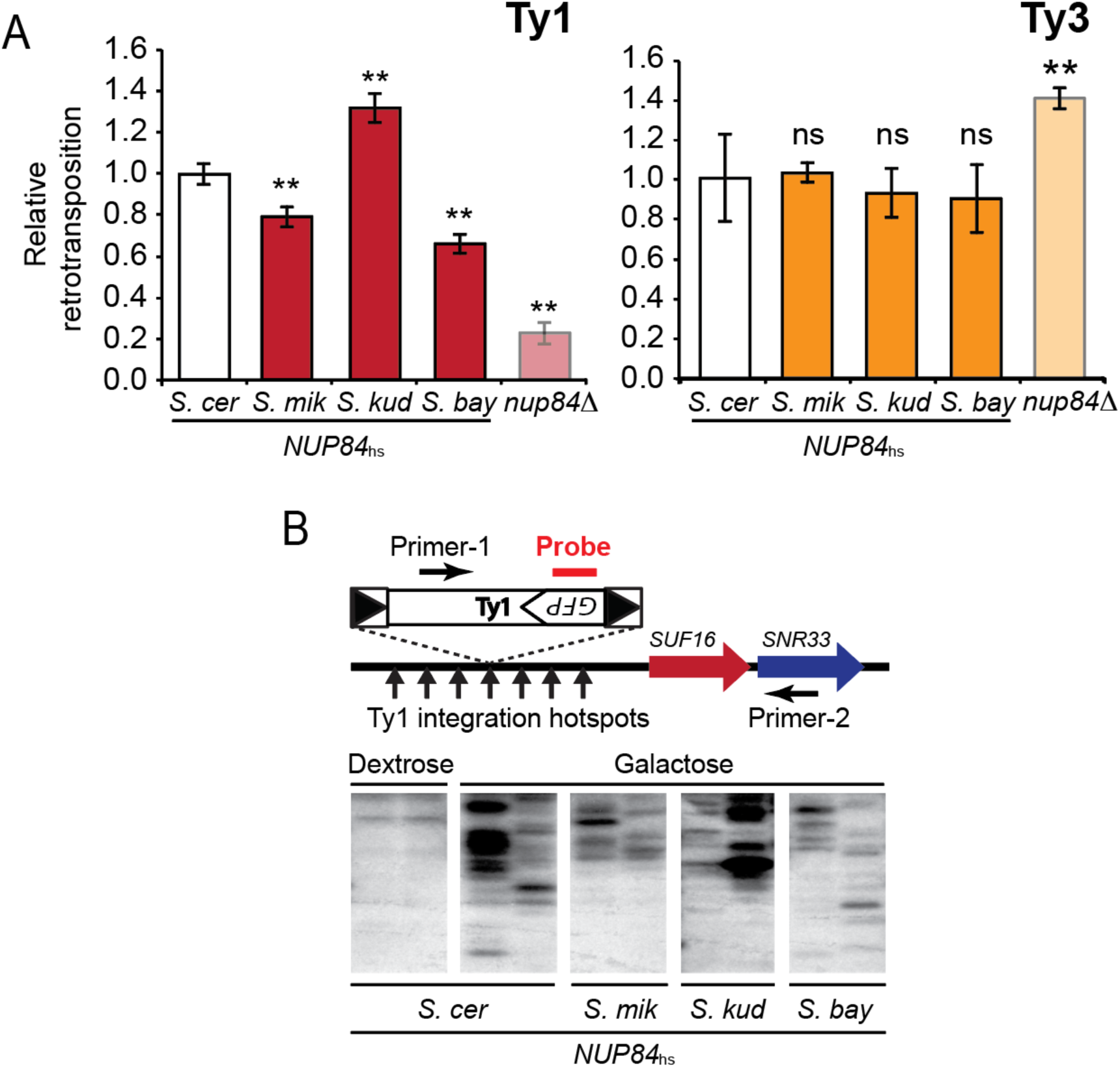
Evolutionary differences between NUP84 of different Saccharomyces species alter levels of Ty1 retrotransposition. (A) Relative retrotransposition of Ty1 and Ty3 within *nup84D* or *nup84D* complemented with heterospecific *NUP84* from different *Saccharomyces* species. Asterisks designate complemented strains that have significantly different levels of retrotransposition compared to the strain encoding *NUP84* from *S. cerevisiae* (Tukey-Kramer method, p<0.05) (error bars: standard error, n>3). (B) Southern blot analysis of Ty1 integration in two independent clones upstream of the *SUF16* locus, which contains Ty1 integration hotspots in its promoter [71]. PCR products across the *SUF16* locus were run on a gel and then probed with a radiolabeled DNA probe specific to *GFP* in order to detect integration events.

These data show that evolutionary differences between *NUP84* of different *Saccharomyces* species modulate the retrotransposition efficiency of Ty1 in a species-specific manner, even though all host functions are conserved. Pairing Ty1 from *S. cerevisiae* with *NUP84* of other species apparently decouples a finely co-evolved relationship, altering levels of retrotransposition. To support this model, we also assayed the impact of *NUP84* evolution on Ty3 replication. We used a galactose inducible Ty3 with a *HIS3* reporter gene and assayed the appearance of colonies able to grow on histidine deficient media as an indication of successful retrotransposition (see methods) [68,72–74]. In contrast to Ty1, we found that *nup84Δ* resulted in increased retrotransposition, as was previously reported [55]. However, each of the heterospecific *NUP84* genes returned transposition to the lower level with no significant difference in retrotransposition among strains encoding heterospecific *NUP84* (one way ANOVA, p=0.90) (Figure 5A). Collectively, these data suggest that the co-evolutionary dynamics are specific to *NUP84* and Ty1.

### *NUP82* has evolved to limit Ty3 retrotransposition

Our evolutionary analysis also identified the gene *NUP82* as being the highest scoring nucleoporin in our evolutionary screen (Figure 2B; 2C; S1), however no role for *NUP82* has been reported in Ty biology. This could be because *NUP82* is an essential gene and would have eluded detection in genome-wide knockout screens. To investigate whether *NUP82* is involved in Ty replication, a dominant negative approach was adopted. Full-or partial-length portions of *NUP82* were expressed in cells that are otherwise wild type at the *NUP82* locus. These Nup82p constructs included the mutations D204A, F290A, Y295A, L393A, I397A, L402A, L405A and F410A (Nup82p^*DFY-LILLF*^) that inactivate interaction with other nucleoporins and decouple it from the nuclear pore complex [75] (Figure 6A). Nup82p^*DFY-LILLF*^ is non-functional as a nucleoporin, therefore we reasoned that it would compete with wild-type Nup82p and have an inhibitory effect on retrotransposition if Ty interacts with Nup82p to transit the nuclear pore. Indeed, the expression of the C-terminal helical domain of Nup82p (residues 433-713) significantly reduced Ty1 retrotransposition, with the N-terminal b-propeller domain (residues 1-458) being dispensable for this effect (Figure 6B). Expression of any of the dominant negative *NUP82* genes did not noticeably affect the growth of *S. cerevisiae* (Figure 6C) or general nuclear import (Figure 6D) compared to expression of the control gene *MET17,* which suggests that these proteins are not toxic to *S. cerevisiae* and do not disrupt the nuclear pore complex. In summary, this serves as preliminary evidence of a previously uncharacterized role for *NUP82* in Ty1 replication.

**Figure 6.**
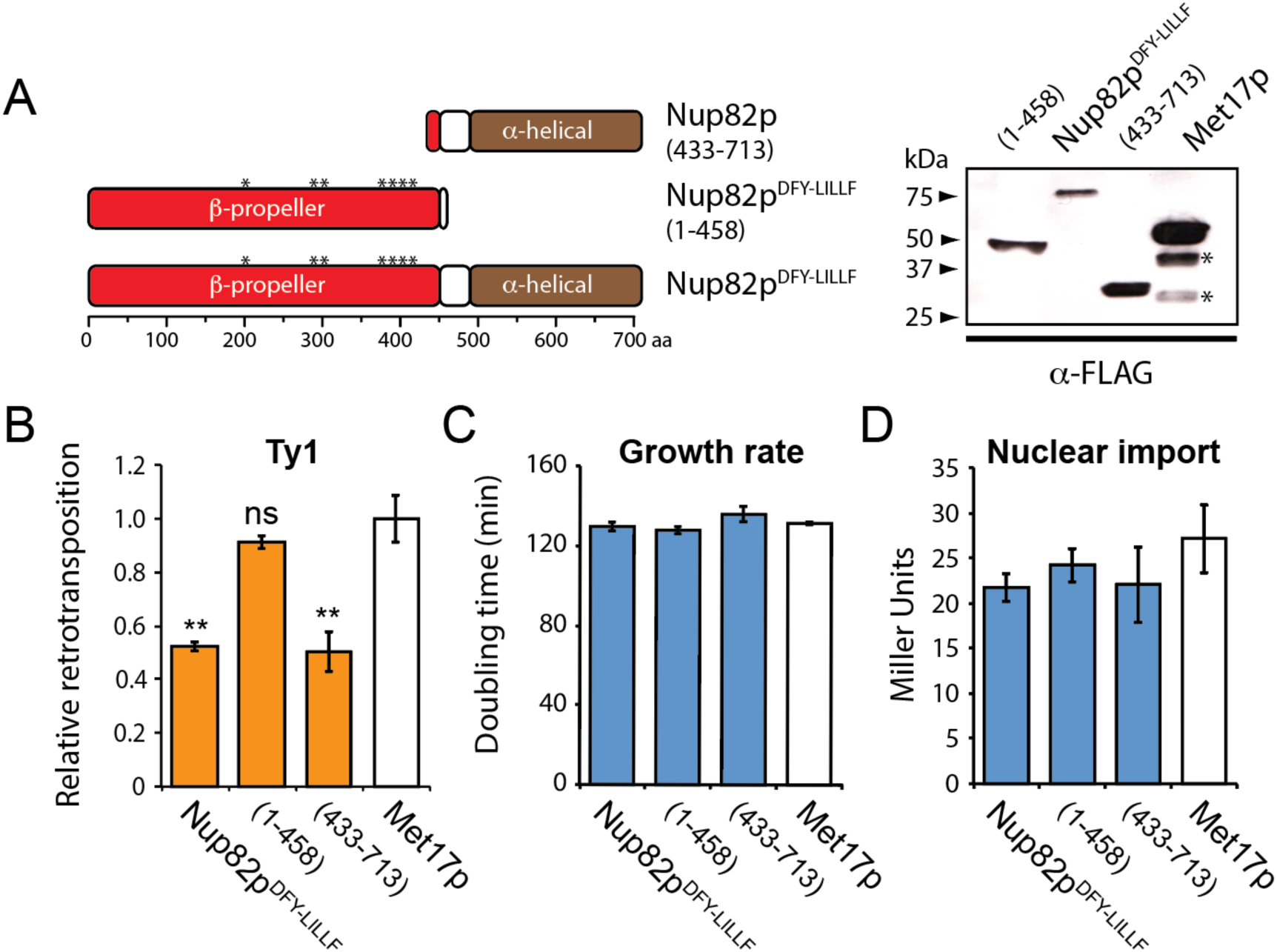
The expression of dominant negative NUP82 and its impact on Ty1 retrotransposition. (A) *Left. A* linear domain diagram of Nup82p^*DFY-LILLF*^ and derived deletion mutants [Nup82p(433-713) and Nup82p^*DFY-LILLF*^(1-458)]. Asterisks mark the mutations that decouple Nup82p from the nuclear pore complex. *Right.* Western blot analysis to detect the expression of FLAG-tagged Nup82p^*DFY-LILLF*^ and its derivatives, compared to the expression of a control protein (Met17p) in the wild-type background (*Met17p degradation products). The effect of Nup82p^*DFY-LILLF*^ expression on (B) Ty1 retrotransposition, (C) doubling time in liquid medium and (D) the nuclear import of the reporter protein LexA-MBP-Gal4(AD), relative to the expression of *MET17* (error bars: standard error, n>3; **Tukey-Kramer method, p<0.05).

In a similar approach to that taken with *NUP84, S. cerevisiae* was engineered to express *NUP82* from different *Saccharomyces* species to ascertain the impact of evolution on Ty retrotransposition. Due to the essential nature of *NUP82,* we used a *NUP82/nup82D* heterozygous diploid strain from the “synthetic genetic array” collection [76] as our starting strain for the genomic replacement of *NUP82* ^*S.cer*^. A customized SceI restriction endonuclease method was used to improve the efficiency of homologous recombination-based gene replacement (see methods) (Figure 7A and S2). *S. cerevisiae* encoding heterospecific *NUP82* have a normal colony morphology, growth rate (Figure 7B), and no difference in *GAL1* and *CUP1* promoter expression (Figure S3), suggesting that the cells are the same in measurable host functions. We tested the effect of *NUP82* evolution on Ty1 retrotransposition using the GFP fluorescence assay and, in contrast to our studies of *NUP84,* found that retrotransposition levels were similar in strains expressing *NUP82* ^*S.mik*^ and *NUP82* ^*Sbay*^ and *NUP82* ^*S.cer*^, but were significantly higher for strains complemented with *NUP82* ^*Skud*^ (Figure 7C). Thus, although *NUP82* may be important for Ty1 retrotransposition (Figure 6), we find that Ty1 seems mostly insensitive to the evolutionary differences between *NUP82* of different species. We next assayed the replication of a Ty3 retrotransposon in the engineered *NUP82* heterospecific strains. *S. cerevisiae* expressing *NUP82* ^*S.mik*^ resulted in a significant >3-fold increase in Ty3 retrotransposition, relative to *NUP82* ^*S.cer*^ (Tukey-Kramer method, p<0.05) (Figure 7C). These data show that the evolutionary differences within *Saccharomyces NUP82* can impact both Ty1 and Ty3 retrotransposition, but predominantly Ty3. Together, we show that *NUP82* appears to play a previously uncharacterized role in Ty retrotransposition, and that Ty1 and Ty3 are differentially susceptible to evolutionary changes within *NUP82*.

**Figure 7.**
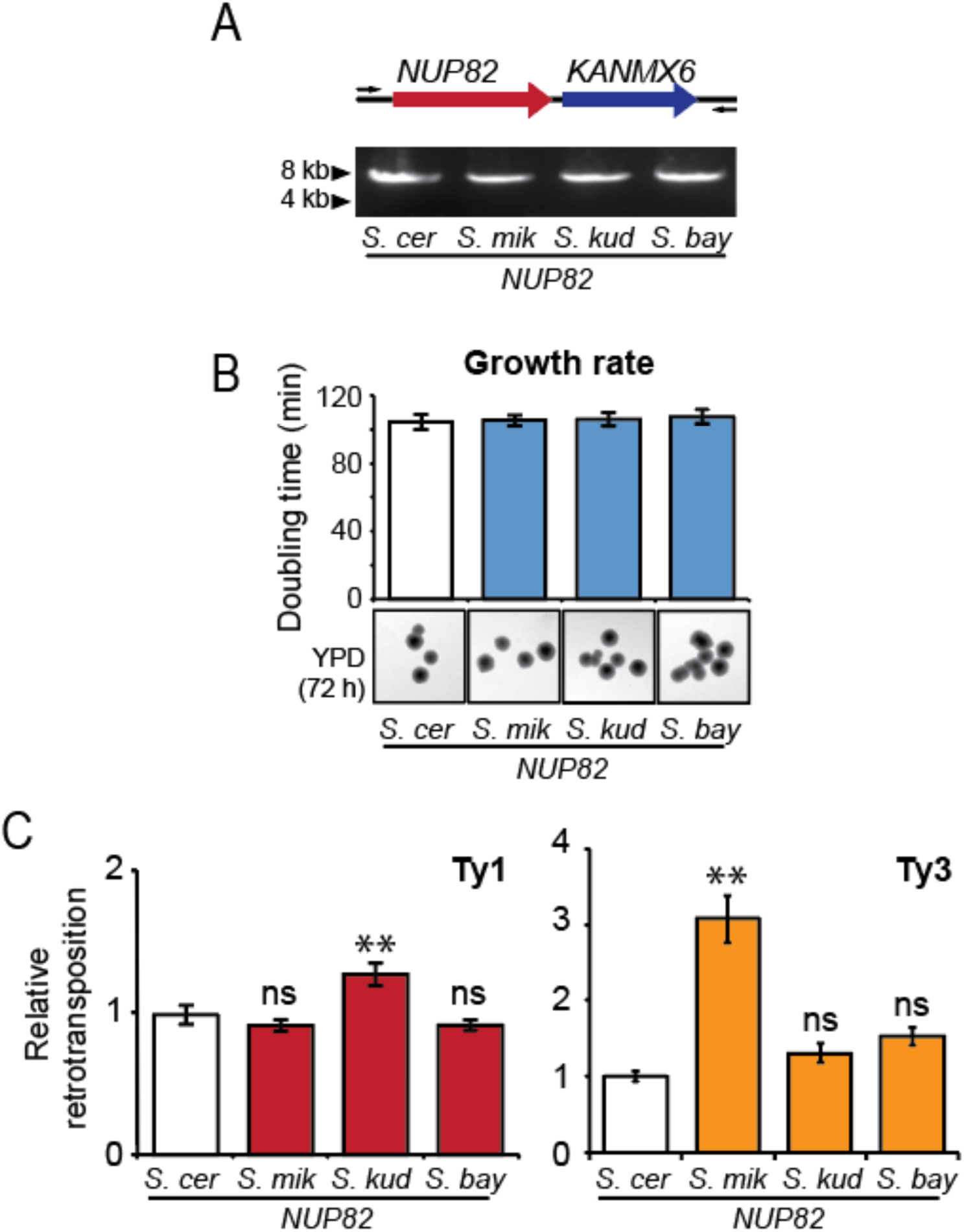
The evolution of NUP82and its effect on Ty1 and Ty3 retrotransposition within *S. cerevisiae*. (A) A schematic representation of *S. cerevisiae* engineered to express heterospecific *NUP82 (NUP82)* from different *Saccharomyces* species. Genome engineering was confirmed by PCR amplification across the *NUP82* locus. (B) The doubling time of *NUP82-* complemented strains in liquid medium. Colony growth and morphology of the engineered strains was monitored for 72 h on solid YPD medium (error bars: standard error, n>3). (C) Relative retrotransposition of Ty1 and Ty3 within strains complemented with *NUP82* from different *Saccharomyces* species. Asterisks mark significant differences in retrotransposition compared to the strain encoding *NUP82* from *S. cerevisiae* (error bars: standard error, n>3; **Tukey-Kramer method, p<0.05).

## DISCUSSION

There are many selective pressures driving the evolution of *Saccharomyces* yeasts, including resource competition, sexual selection, and long-term co-evolution to exist with viruses and other genetic parasites [11,12,77–81]. As Tys are the only known genetic parasites within *Saccharomyces* yeasts that enter the nucleus, this provides a unique opportunity to study their impact on the evolution of the nuclear pore complex. We find signatures of natural selection acting on several nucleoporins, coinciding with previous observations that deletion or disruption of several of these genes alters Ty retrotransposition. Here, we use a unique approach to demonstrate that the evolutionary changes that have accumulated in yeast nucleoporins alter Ty retrotransposition levels. We replaced *NUP82* and *NUP84,* within the context of the *S. cerevisiae* genome, with orthologs from related *Saccharomyces* yeasts, and then measured Ty retrotransposition in these isogenic yeast strains. It is important to note that the genetic parasites have been held constant in the study, with both Ty1 and Ty3 deriving from the *S. cerevisiae* lineage. In some cases, orthologs of *NUP82* and *NUP84* resulted in higher levels of *S. cerevisiae* Ty retrotransposition, and in other cases, they resulted in lower levels. These patterns are consistent with a model where nucleoporins are co-evolved with the Ty of their own species. When a Nup82p or Nup84p ortholog is substituted within the *S. cerevisiae* nuclear pore, sometimes *S. cerevisiae* Ty can exploit it better than it can the *S. cerevisiae* version of that protein (possibly by having an increased affinity for the foreign ortholog, which is not evolved to evade it). Other times, the *S. cerevisiae* Ty is less able to utilize this orthologous protein because the Ty has not evolved refined interaction with this particular ortholog. Ultimately, our data shows that uncoupling *S. cerevisiae* Tys from the co-evolved *S. cerevisiae NUP82* or *NUP84* results in altered levels of retrotransposition. The replacement of *S. cerevisiae* nucleoporins with heterospecific nucleoporins “decouples” this evolutionarily optimized interaction and leads to either an increase or decrease in retrotransposition, without impacting cellular homeostasis (e.g. nuclear import). While it is never possible to know for sure what has driven selection within these genes, nucleoporins from different *Saccharomyces* species support variable levels of Ty1 or Ty3 retrotransposition, providing a phenotypic trait on which selection may have been acting. This is similar to our recent observations that the antiviral *XRN1* gene from *Saccharomyces* yeasts has likely co-evolved with totiviruses to control excessive viral replication [11].

The exact functions of *NUP82* and *NUP84* during Ty retrotransposition, and their mechanism of action, remain unclear. Ty nuclear ingress likely involves docking of the virus-like particle to the nuclear periphery by interaction with nucleoporins [25,41]. The known positioning of Nup82p and Nup84p at the cytoplasmic face of the nuclear pore complex could possibly facilitate virus-like particle docking, in a similar manner to their recruitment and binding of host karyopherins prior to nuclear import [82–85]. Multiple Ty3 proteins (Gag3, p27 and CA) interact directly with nucleoporins, and the integrase of Ty1 and Ty3 contain nuclear localization signals [25,42–45]. Therefore, it seems likely that Ty proteins interact directly with nucleoporins, and that evolutionary selection could be acting to alter these physical interactions. That fragments of Nup82p inhibit retrotransposition is consistent with this. Because these fragments aren’t incorporated into the nuclear pore, they are likely acting through dominant negative physical interactions with Ty.

The evolutionary relationship between yeast and Ty retrotransposons is complex. The intracellular lifecycle and ubiquity of Ty would suggest that Ty have been co-evolving with the *Saccharomyces* genus for many millions of years [1,2]. Evolutionary selection would favor limited Ty replication, as active retrotransposition and high Ty copy number can alter yeast fitness [3–6,10]. Indeed, certain families of Tys are absent from certain strains and species of *Saccharomyces* yeasts [1,64,86,87]. However, the persistence of Tys in *Saccharomyces* yeasts suggests that loss of Tys is rare, perhaps due to Ty introgression and transmission by sexual reproduction, which are potential mechanisms by which Tys can invade Ty-free or naive populations [1,86,88]. Also, the error prone nature of the Ty reverse transcriptase and reverse transcription-mediated recombination can generate Ty variants that could also overcome host-encoded resistance mechanisms [89]. In contrast to the idea that Tys are completely parasitic, Ty retrotransposition can drive the evolution of the yeast genome by changing gene regulation and expression by integrating in or near host genes. Tys can also facilitate gross chromosomal rearrangements of the host genome, including translocations and deletions, by way of homologous recombination between Ty integrated at different locations within host chromosomes [90–93]. Experimental systems have shown that Ty-mediated genome evolution can be observed in the laboratory [4], and would likely allow populations of *Saccharomyces* yeasts to rapidly respond to selective pressures found within the natural environment. Thus, in the context of the nuclear pore complex, there may be evolutionary selection to prevent Ty nuclear transit and excessive replication, but also selection against mutations that completely abrogate retrotransposition. The long-term association between *S. cerevisiae* and its cognate Tys would imply that this interaction has been “optimized” by evolutionary selection, perhaps to balance the damaging effects of excessive Ty retrotransposition with the benefits of genome plasticity.

The nuclear pore is the gatekeeper of the nucleus, and it is antagonized by many pathogens and genetic parasites. The nucleoporins that are under positive selection in yeast have also been shown to be essential to the replication of other retrotransposons and viruses. The fission yeast *Schizosaccharomyces pombe* ortholog of *NUP1* (*NUP124*) is required for retrotransposition of the Ty3/gypsy-like element Tf1 and directly interacts with the Tf1-encoded Gag protein [94,95]. The human homologs of *NUP1* and *NUP116 (NUP153* and *NUP98,* respectively) are important for viral replication in humans, including for HIV, HBV, HCV, and influenza virus [24,29,33,96–101]. Specifically, *NUP153* is an important determinant of HIV and HBV nuclear import, and its FG (phenylalanine-glycine)-repeat domain directly interacts with HIV capsid, via specific FG-repeats [24,33,102,103]. In *S. cerevisiae,* the FG-repeat region of Nup116p directly interacts *in vitro* with the Ty3-encoded protein Gag3, and truncation of *NUP116* decreases retrotransposition [25]. In human cells, the reduced expression of *NUP88* and *NUP107* (orthologs of yeast *NUP82* and *NUP84,* respectively) reduces HIV and influenza virus replication [97,104], however, it is unclear whether their role in viral nuclear import is direct or a consequence of pleiotropic effects (e.g. disruption of the nuclear pore complex). Collectively, this paints a picture of complex evolutionary pressures on nuclear pore genes across eukaryotes.

It appears that viral infections have broadly shaped the evolution of host genomes, affecting genes well beyond immunity loci [105]. The most classic example involves cellular entry receptors used by viruses to enter cells. These receptors are often under positive selection, resulting in highly species-specific interactions with viruses [106–110]. The nuclear pore complex is the gatekeeper of the nucleus just like cell surface receptors are gatekeepers of the cytoplasm. Our work in *Saccharomyces* yeasts provides a framework to further investigate the importance of the nuclear pore complex in modulating Ty retrotransposition, and for a parallel investigation into the evolution of the orthologous nuclear pore complex of higher eukaryotes. It remains unknown how broadly viruses and genetic parasites are driving the evolution of important housekeeping proteins, but intriguing recent reports involving genes such as *XRN1* (involved in degradation of uncapped mRNAs; [11]), and DNA repair genes [111], suggest that this might be more common than previously appreciated.

## Materials and Methods

### Plasmid construction

The *ACT1* intron (ACT1i) and an artificial intron (AI) [112] were amplified by PCR with included primer-encoded flanking homology to *GFP.* This PCR product was inserted directly after the ATG start codon at the 5’ end of *GFP* by the “yeast plasmid construction by homologous recombination” method (recombineering) [113]. GFP(AI) and GFP(ACT1i) were amplified by PCR and introduced into pAG423-GAL-ccdB using TOPO-TA and Gateway cloning strategies (Thermo Fisher) to create pPAR061 and pPAR063, respectively. GFP(ACT1i) was also placed under the control of the *CUP1* inducible promoter (456 bp upstream of *CUP1* were cloned directly from the genome of S.*cerevisiae)* using recombineering. pCUP1-GFP(ACT1i) was used to replace H/S3(AI) within pGTy1-HIS3(AI) to create pPAR078. pPAR101, pPAR104, pPAR145 and pPAR181 were constructed by using PCR to create DNA encoding FLAG-tagged Nup82p^*DFY-LILLF*^(1-458), Nup82p^*DFY-LILLF*^ and Nup82p (433-713) from pNOP-GFP-Nup82p^*DFY-LILLF*^ [75]. *MET17 was* amplified directly from the genome of *S. cerevisiae.* All PCR fragments were subsequently cloned into pAG414-GPD-ccdB via pCR8 using TOPO-TA and Gateway cloning strategies (Thermo Fisher). To assay nuclear import using the strategy outlined by Marshall *et al.* we first subcloned the LexA-MBP-GAL4(AD) cassette from pJMB1076n [69] into the pAG413 plasmid backbone using recombineering, essentially changing the selective marker on the plamsid from *LEU2* to *HIS3.* For gene knockout, all plasmids were constructed by recombineering using *NUP82* and *NUP84* amplified from various *Saccharomyces* species. These nucleoporin genes were placed upstream of a selectable marker *(LEU2* or *KANMX6)* and the entire cassette flanked by 1000 bp of sequence encompassing the 5’ and 3’ untranslated regions of *NUP82* or *NUP84* from *S. cerevisiae.* pPAR240 was constructed by first amplifying a LexA operator sequence upstream of the p-galactosidase gene from *S. cerevisiae* L40. PCR products were designed to contain flanking homology to *ADE2* from *S. cerevisiae* and these PCR products were used to disrupt the *ADE2* gene within pRS422 to create pPAR240.

### Evolutionary analysis

Gene sequences from six species of *Saccharomyces* yeasts were obtained from publically available online resources. Maximum likelihood analysis of dN/dS was performed using the codeml program in PAML 4.1. Multiple protein sequence alignments were created and were manually curated to remove ambiguities before processing with PAL2NAL to produce accurate DNA alignments [114]. DNA alignments were fit to two models: M7 (codons fit to a beta distribution of dN/dS values, with dN/dS > 1 disallowed) and M8 (similar to model 7, but with dN/dS > 1 allowed). Two models of codon frequencies (f61 and f3x4) and multiple seed values for dN/dS (w) were used (File S1). Likelihood ratio tests were performed to evaluate which model of evolution the data fit significantly better with positive selection and inferred if we can reject M7 in favor of M8 with a p<0.05. REL and FEL codon based models were also used to detect sites under positive selection as implemented by the HyPhy package using the best substitution models chosen by Akaike information criterion (AIC) using the phylogenetic tree (Newick format): ((((*S. paradoxus, S. cerevisiae), S. mikatae), S. kudriavzevii), S. arboricolus, S. bayanus*).

### Strain Construction

Standard methodologies for PCR-based gene knockout and replacement were used to create all *NUP84* strains in BY4741 (YPAR0130-0133) [115]. Strains YPAR0135-0138 were engineered to encode a LexA operator sequence upstream of the p-galactosidase gene, and were constructed by the disruption of the genomic copy of *ADE2* using a PCR cassette amplified from pPAR240. Clones selected for their ability to grow on media lacking uracil and inability to grown on media lacking adenine. *NUP82* gene replacement utilized a SceI-based method to increase the efficiency of the integration of *NUP82* and *KANMX6* by generating DNA double-stranded breaks at the *NUP82* locus in *S. cerevisiae* (personal communication, Dr. C.M. Yellman). Using a diploid heterozygous knockout strain of *NUP82* [76], *KANMX6* at the *NUP82* locus was replaced with the *URA3* gene from *K. lactis* flanked by SceI sites amplified by PCR from pCMY-IT3. Gene replacement was carried out by the concomitant expression of SceI from pGAL1-SCEH and the LiAc transformation of a PCR-derived cassette encoding *NUP82-KANMX6. NUP82/nup82D::NUP82-KANMX6* clones were selected by their ability to grow in the presence of 400 μg mL^−1^ G418 and their resistance to 5-FOA (0.1% w/v). Haploid clones were isolated from the engineered diploid strains using the SGA selection protocol as described previously [76]. The correct insertions were confirmed by PCR of genomic DNA of the *NUP82* locus to create strains YPAR0139, YPAR0143, YPAR0141 and YPAR0142. A PCR cassette was used to disrupt *HIS3* in YPAR0139, YPAR0143, YPAR0141 and YPAR0142, clones were selected for their ability to grow on media containing hygromycin and inability to grown on media lacking histidine to produce strains YPAR0143, YPAR0145, YPAR0147 and YPAR0149.

### Splicing of the ACT1 intron and insertion of an artificial intron within the GFP gene

Plasmids pPAR063 and pPAR061 were used to produce *GFP* transcripts containing either the *ACT1* intron (ACT1i) or an artificial intron (AI) [112], respectively, by induction from a galactose inducible promoter. Cultures were grown to mid-log phase in liquid culture with raffinose as the sole carbon source. At OD_600_ of ~1 galactose or dextrose were added to a final concentration of 2% and the cultures grow at 30°C for 2 h. Total RNA was extracted from these cultures (~2 × 10^7^ cells) using the RNeasy RNA extraction kit (Qiagen). 5 μg of RNA was treated with 1 U of DNase I at 37°C for 10 min before heat inactivation at 7°C for 10 min. RNA samples were then subject to two-step RT-PCR using the Superscript III one-step method using the GFP-specific primers 5’-AAGCTGACCCTGAAGTTCATCTGC-3’ and 5’-CGTTGTGGCTGTTGTAGTTGTACTCC-3’.

### Ty1 retrotransposition assays

Yeast strains to be assayed for their ability to support Ty1 retrotransposition were transformed with either pPAR078 (GFP flow cytometry method) or pGTy1(HIS3(AI)) (as previously described in [71]). To detect retrotransposition using GFP positive cells, single colonies from *S. cerevisiae* transformed with pPAR078 were isolated for each new experiment. Each experiment was performed at least three times. Colonies were first grown for 24 h in 2 mL raffinose-uracil complete medium at 30°C with agitation. 1 × 10^5^ cells from the saturated cultures were used to inoculate 15 mL of complete medium with galactose-uracil, followed by growth for 5 days at room temperature with agitation. Cultures were diluted and allowed to reach early log phase growth (OD_600_ ~0.05) before the addition of CuSO_4_ to a final concentration of 0.5 mM. Cultures were grown for 9 h at 30°C before assaying for the presence of live, GFP positive cells using a BD LSRII Fortessa flow cytometer (San Jose, CA) running FACSDiva software (v6.1.3). GFP excitation was observed with a blue, 488nm laser, while GFP emission was collected using 530/30nm band pass filter and 502nm long pass filter. Propidium iodide (PI) excitation was observed with a yellow-green, 561nm laser, while PI emission was collected using 660/20nm band pass filter and a 635nm long pass filter. 100,000 gated events were collected using a forward scatter vs. side scatter dot plot, with forward scatter showing relative particle size and side scatter showing internal complexity. All subsequent plots were generated from this gated population. Live cells were gated by staining cell populations with PI (final concentration 0.1 μg mL^−1^) and GFP positive populations were gated by comparison with GFP negative populations of cells. Analysis of flow cytometry data was performed using FlowJo version 9.7.6.

### Ty3 transposition assay

For quantification of Ty3 transposition, yeast cells were transformed with pPS3858, a *URA3* marked galactose inducible Ty3-HIS3 [68,72–74]. The *HIS3* gene is located at the end of *POL* and is anti-sense to Ty3, except for an artificial intron which is sense. The sense intron prevents production of His3p until after the full-length Ty3 RNA is transcribed, spliced, reverse transcribed and integrated into the genome. Colony transformants were selected on synthetic media with 2% glucose (SD) complete with amino acids but lacking uracil. Single colonies were inoculated into 2 mL of synthetic raffinose (-uracil) and grown for 24 h. Cultures were then brought to 5 mL and grown for ~8 h, after which OD_600_ was measured and cultures were diluted back to an OD_600_ of ~0.02 in 4.5 mL and grown overnight. The following morning, 500 μL of 20% galactose (2% final) was added to induce Ty3 expression; after 8 h of induction cultures were pelleted, washed in SD media, serially diluted, and plated on both SD plates lacking histidine for growth of transposed cells and also YPD plates to determine total live cell counts.

### Nup82p^*DFYLILLF*^ expression assays

Plasmids constitutively expressing either Nup82p^*DFY-LILLF*^, Nup82p truncation mutants or Met17p were introduced into a strain containing a Ty1 retrotransposition reporter plasmid. Retrotransposition assays were carried out as previously described above, but with the use of double-dropout complete medium to maintain both episomal vectors during retrotransposition.

### Western blotting of Nup82p ^*DFY-LILLF*^ and truncation mutants

Yeast lysates were prepared from 5 mL of stationary phase culture grown for 16 h in yeast complete medium-leucine-tryptophan. Cell pellets were washed with 1 mL of chilled 25 mM Tris-HCl (pH 7.0), 10 mM sodium azide before incubation at 100°C for 3 min. 50 μL of SDS sample loading buffer (100 mM Tris-HCl, 5% SDS, 10% glycerol, 0.1% bromophenol blue, 2% p-mercaptoethanol, pH 6.8) was added to the boiled pellet with 200 μL of acid-washed glass beads (0.5 mm). Samples were vortexed for 10 min to disrupt yeast cells before the addition of another 80 μL of SDS sample loading buffer. Glass beads were pelleted by centrifugation (1500 × g, 2 min). 30 μL of each sample was loaded directly onto a precast Tris-glycine 10% SDS-PAGE gel (Biorad). Flag-tagged *NUP82* mutants were detected via Western blot using a 1:4000 dilution of a primary mouse monoclonal anti-flag (Syd Labs #M20008). Secondary detection was carried out using a 1:2000 dilution of a goat anti-mouse horseradish peroxidase conjugated antibody (Thermo #32430).

### Nuclear pore complex import assays

A LexA-MBP-GAL4(AD) fusion protein with or without an SV40 nuclear localization signal [69] was used to measure the efficiency of nuclear import within *S. cerevisiae* L40 or BY4741. 5 mL of glucose supplemented synthetic complete medium lacking the appropriate amino acid and grown overnight at 30°C with agitation. Cells were collected by centrifugation at 4000 × g for 30 seconds and the cell pellets suspended in 750 μL of ice-cold ddH_2_O. Washed cells were again collected by centrifugation (13,000 x g for 30 seconds), and soluble proteins extracted by Y-PER buffer as per manufacturer’s instructions (Thermo). The lysate was assayed for μ-galactosidase activity as described previously [116].

### Fluorescence microscopy

The steady-state import of GFP-NLS was monitored within BY4741 transformed with pEB0836 as described previously [70].

### Detection of Ty1 genomic integrations by Southern blotting

The detection of the integration of Ty1 containing *GFP* by Southern blotting was performed as previously described [71], in the various NUP84-complemented or deletion strains of *S. cerevisiae.* Total DNA was extracted from cell cultures using phenol:chloroform and ethanol precipitation, after 5 days of retrotransposition induction, as described in the Ty1 retrotransposition assay protocol above. Southern blotting was carried out after agarose gel electrophoresis, as described previously [117], using Hybond-XL membranes (GE healthcare).

### Promoter activity assay

To assay the activity of the *GAL1* promoter we expressed *GFP* under the control of the *GAL1* promoter and monitored the increase in the mean fluorescent intensity (MFI) compared to uninduced control cells. Cells were grown overnight to saturation at 30°C (CM-uracil, 2% raffinose) before being used to seed a 10 mL culture that was grown to log phase (OD_600_ 0.1-0.5). Each 10 mL log phase culture was divided into two 5 mL cultures, supplemented with either 2% galactose or dextrose (final concentration) and grown for 6 h. Cultures were assayed for GFP fluorescence by flow cytometry using the same instrumentation as described above. The activity of the *CUP1* promoter was assayed by analyzing MFI data derived from Ty1-GFP retrotransposition assays.

## Acknowledgments

The authors would like to thank Emily Feldman, David Garfinkel, Chien Hui-Ma, André Hoelz, Makkuni Jayaram, Aashiq Kachroo, Maryska Kaczmarek, Nicholas Meyerson, Joe Mymryk, Soumitra Sau, Matthew Sorenson, Alex Stabell, Scott Stevens, Cody Warren, Suzanne Wente, Chris Yellman and Renate van Zandwijk for critical reagents, laboratory support, and insightful discussions. This work was supported by a grant from the National Institutes of Health (GM093086 to S.L.S.). S.L.S. is a Burroughs Wellcome Fund Investigator in the Pathogenesis of Infectious Disease.

## Supporting Information

### Supplementary Figure Legend

**Figure S1.**
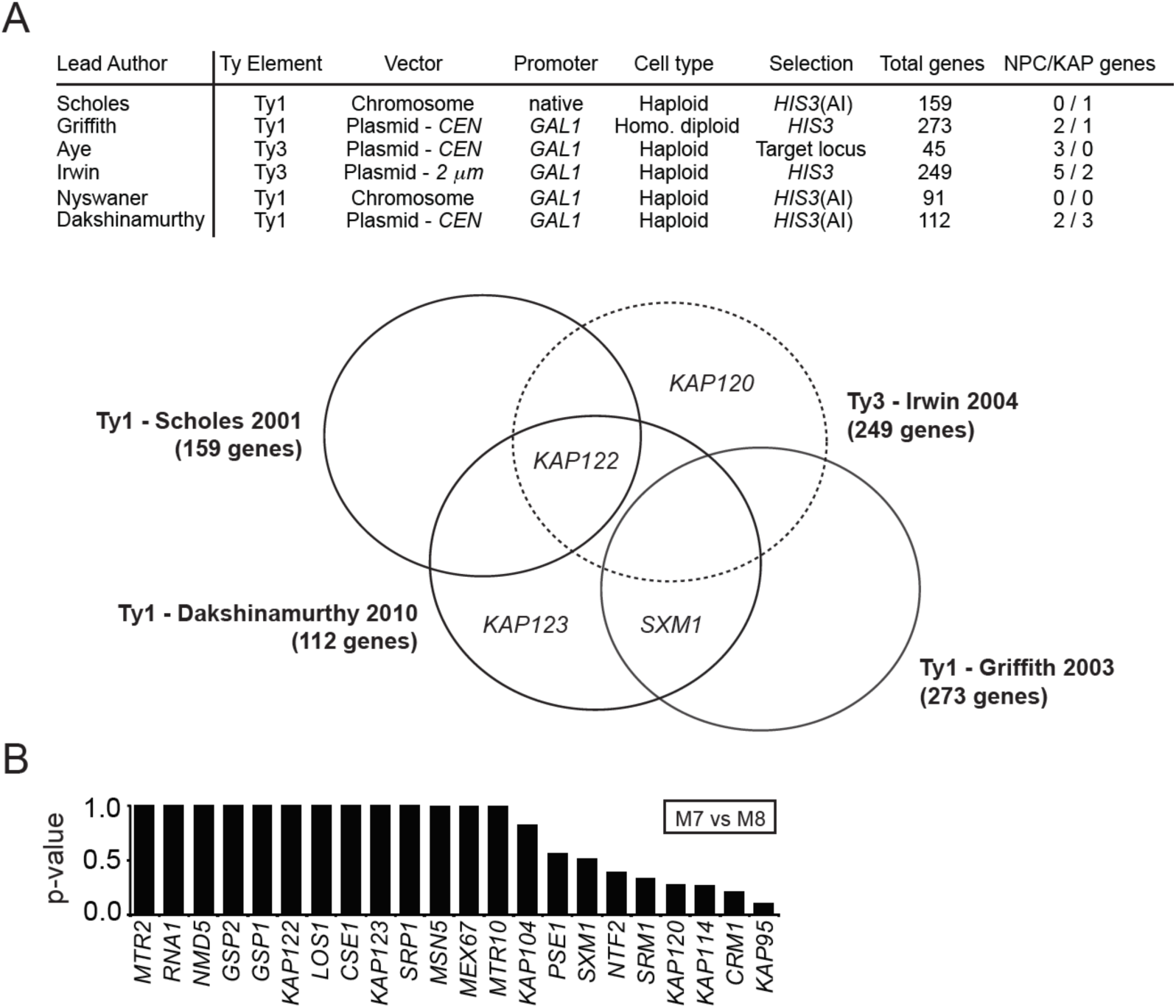
Nucleoporins and karyopherins are important for Ty retrotransposition, but karyopherins are not evolving rapidly. (A) A summary of whole genome studies that have identified nucleoporins and karyopherins important for Ty1 and Ty3 retrotransposition [51–56]. (B) Results from PAML analysis surveying karyopherins for signatures of positive selection, comparing a codon model of purifying selection (M7) to a codon model of positive selection (M8). No karyopherins had a p<0.05.

**Figure S2.**
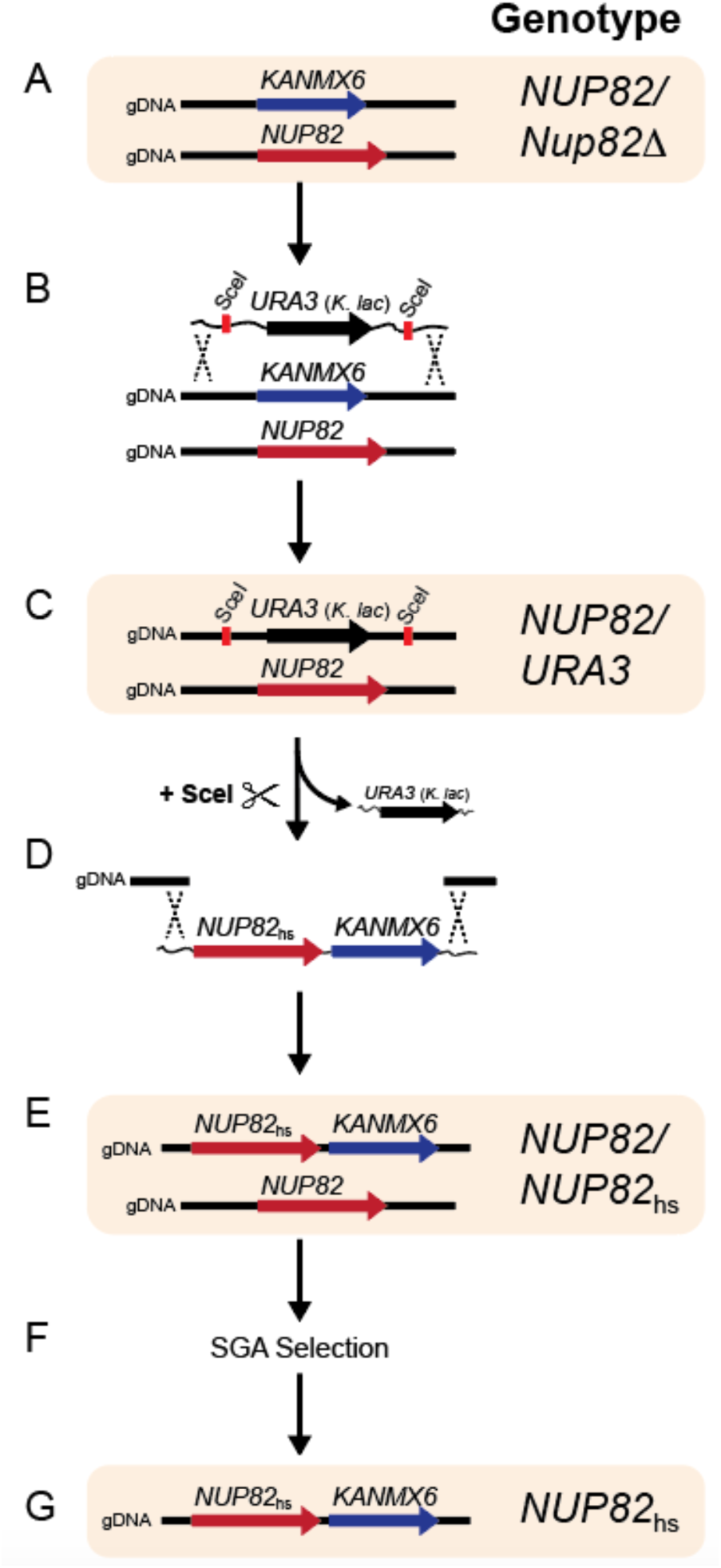
The construction of *S. cerevisiae* strains expressing heterospecific *NUP82*. The *KANMX6* gene within a diploid strain of *S. cerevisiae* heterozygous for *KANMX6* at one *NUP82* locus (A) was replaced with the *URA3* gene from *K. lactis* flanked by SceI sites (B-C). SceI restriction endonuclease was used to create double-stranded DNA breaks at the URA3-containing *NUP82* locus, which was simultaneously repaired by a PCR-derived cassette encoding heterospecific *NUP82* and *KANMX6* (D-E). Haploid clones were isolated using the SGA selection protocol [76] (F-G).

**Figure S3.**
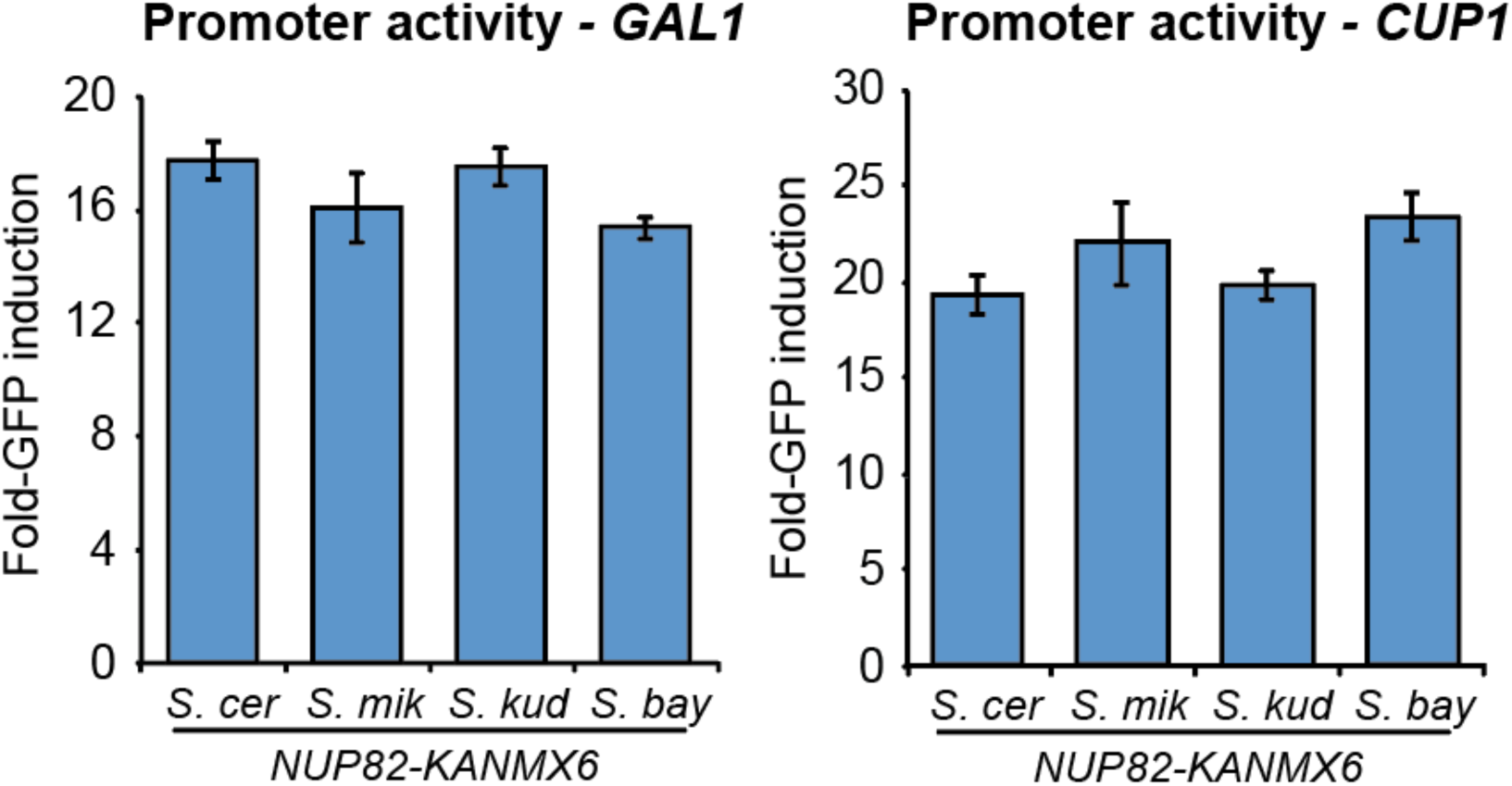
The evolution of *NUP82* does not impact GFP production from different promoters. The effect of *NUP84* complementation on the ability of *S. cerevisiae* to express GFP from the promoters used in our Ty1 GFP-based reporter *(GAL1* or *CUP1* promoters).

**Table S1.**
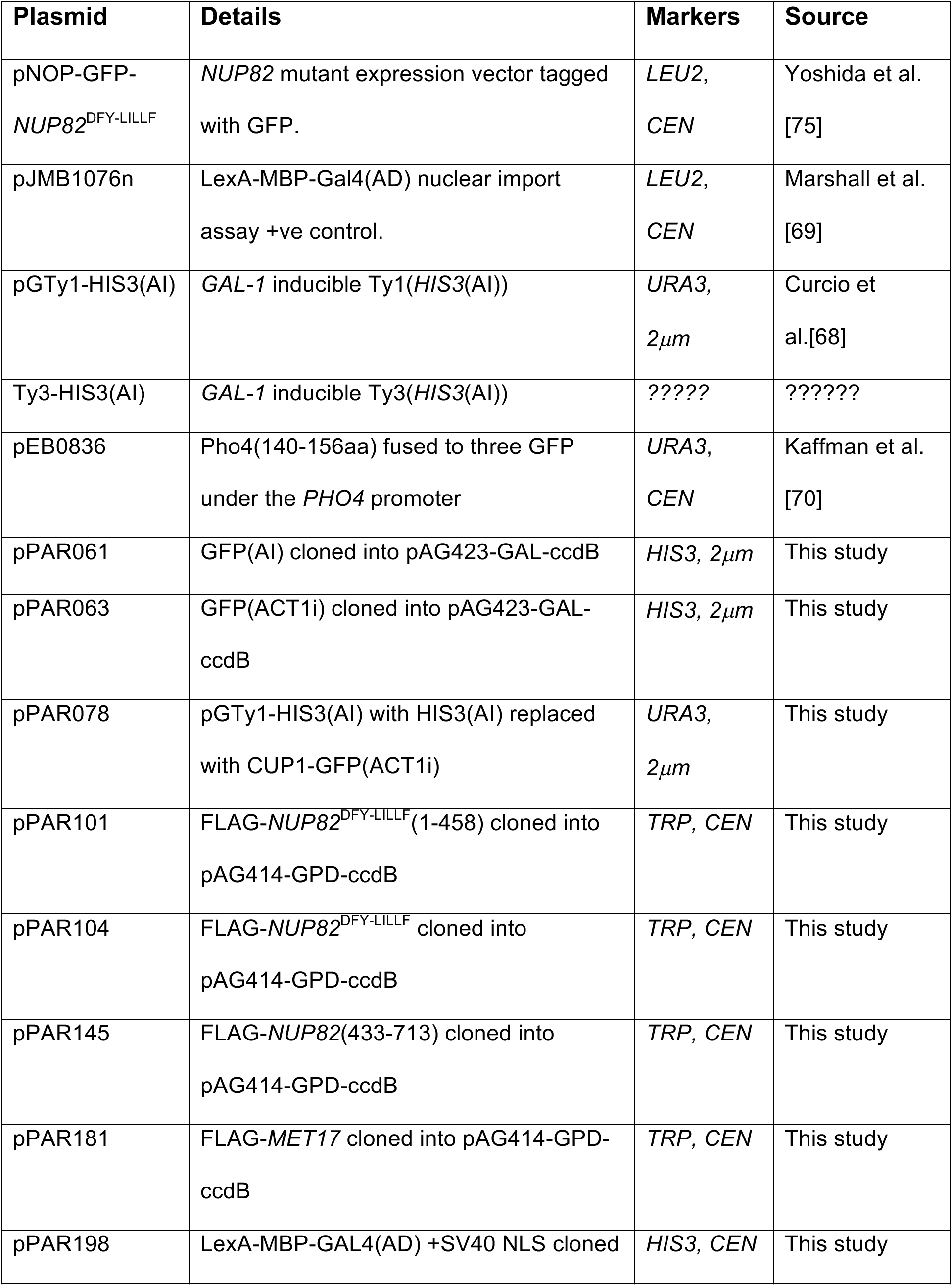
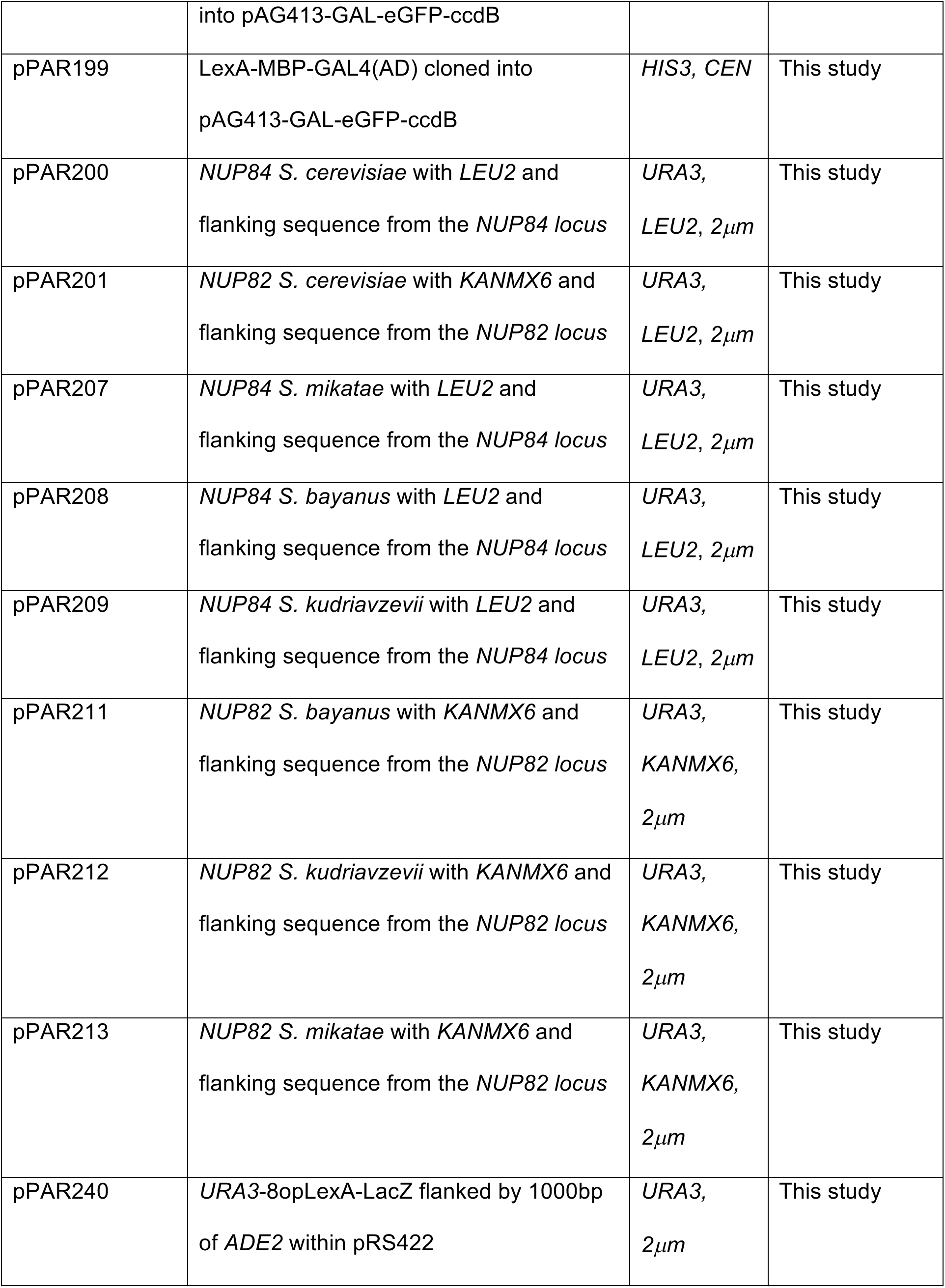
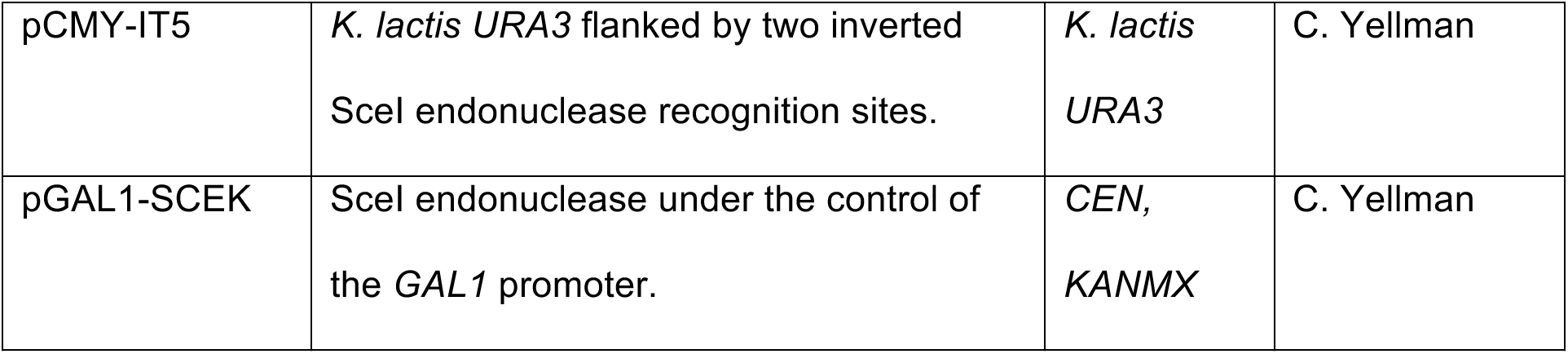
Plasmids

**Table S2.** Yeast strains

| **Strain** | **Genotype** | **Source** |
| --- | --- | --- |
| BY4741 | MATa *his3Δ1 leu2Δ0 met15Δ0 ura3Δ0* | [118] |
| BY4743 *nup82Δ/NUP82* | MATa/α *ura3Δ0 leu2Δ0 his3Δ1*<br>*LYS2+/lys2Δ0 met15Δ0/MET15+*<br>*can1Δ::LEU2+-MFA1pr-HIS3/CAN1+*<br>*nup82Δ::KANMX/NUP82* | SGA KO collection<br>[76] |
| BY4741 *xrn1Δ* | MATa *his3Δ1 leu2Δ0 met15Δ0 ura3Δ0*<br>*xrn1Δ* | Haploid KO<br>collection [119] |
| BY4741 *nup84Δ* | MATa *his3Δ1 leu2Δ0 met15Δ0 ura3Δ0*<br>*nup84Δ* | Haploid KO<br>collection [119] |
| BY4741 *nup133Δ* | MATa *his3Δ1 leu2Δ0 met15Δ0 ura3Δ0*<br>*nup133Δ* | Haploid KO<br>collection [119] |
| BY4741 *bud22Δ* | MATa *his3Δ1 leu2Δ0 met15Δ0 ura3Δ0*<br>*bud22Δ* | Haploid KO<br>collection [119] |
| BY4741 *xrs2Δ* | MATa *his3Δ1 leu2Δ0 met15Δ0 ura3Δ0*<br>*xrs2Δ* | Haploid KO<br>collection [119] |
| BY4741 *nup100Δ* | MATa *his3Δ1 leu2Δ0 met15Δ0 ura3Δ0*<br>*nup100Δ* | Haploid KO<br>collection [119] |
| BY4741 *nup84Δ* | MATa *his3Δ1 leu2Δ0 met15Δ0 ura3Δ0*<br>*nup84Δ* | Haploid KO<br>collection [119] |
| YPAR0130 | MATa *his3Δ1 leu2Δ0 met15Δ0 ura3Δ0*<br>*nup84Δ::NUP84-LEU2 (S. cerevisiae)* | This study |
| YPAR0131 | MATa *his3Δ1 leu2Δ0 met15Δ0 ura3Δ0* | This study |
|  | *nup84Δ::NUP84-LEU2 (S. mikatae)* |  |
| YPAR0132 | MATa *his3Δ1 leu2Δ0 met15Δ0 ura3Δ0*<br>*nup84Δ::NUP84-LEU2 (S. bayanus)* | This study |
| YPAR0133 | MATa *his3Δ1 leu2Δ0 met15Δ0 ura3Δ0*<br>*nup84Δ::NUP84-LEU2 (S. kudriavzevii)* | This study |
| YPAR0135 | MATa *his3Δ1 leu2Δ0 met15Δ0 ura3Δ0*<br>*nup84Δ::NUP84-LEU2 (S. cerevisiae)*<br>*ade2::URA3::(LexAop)8-LacZ* | This study |
| YPAR0137 | MATa *his3Δ1 leu2Δ0 met15Δ0 ura3Δ0*<br>*nup84Δ::NUP84-LEU2 (S. mikatae)*<br>*ade2::URA3::(LexAop)8-LacZ* | This study |
| YPAR0136 | MATa *his3Δ1 leu2Δ0 met15Δ0 ura3Δ0*<br>*nup84Δ::NUP84-LEU2 (S. bayanus)*<br>*ade2::URA3::(LexAop)8-LacZ* | This study |
| YPAR0138 | MATa *his3Δ1 leu2Δ0 met15Δ0 ura3Δ0*<br>*nup84Δ::NUP84-LEU2 (S. kudriavzevii)*<br>*ade2::URA3::(LexAop)8-LacZ* | This study |
| YPAR0139 | MATa *ura3Δ0 leu2Δ0 his3Δ1*<br>*can1Δ::LEU2+-MFA1pr-HIS3*<br>*nup82Δ::NUP82 (S. cerevisiae)-*<br>*KANMX6* | This study |
| YPAR0143 | MATa *ura3Δ0 leu2Δ0 his3Δ1*<br>*can1Δ::LEU2+-MFA1pr-HIS3*<br>*nup82Δ::NUP82 (S. mikatae)-* | This study |
|  | *KANMX6* |  |
| YPAR0141 | *MATa ura3Δ0 leu2Δ0 his3Δ1*<br>*can1Δ::LEU2+-MFA1pr-HIS3*<br>*nup82Δ::NUP82 (S. kudriavzevii)-*<br>*KANMX6* | This study |
| YPAR0142 | *MATa ura3Δ0 leu2Δ0 his3Δ1*<br>*can1Δ::LEU2+-MFA1pr-HIS3*<br>*nup82Δ::NUP82 (S. bayanus)-*<br>*KANMX6* | This study |
| YPAR0143 | *MATa ura3Δ0 leu2Δ0 his3Δ1*<br>*can1Δ::LEU2+-MFA1pr-his3Δ::HPHX6*<br>*nup82Δ::NUP82 (S. cerevisiae)-*<br>*KANMX6* | This study |
| YPAR0145 | *MATa ura3Δ0 leu2Δ0 his3Δ1*<br>*can1Δ::LEU2+-MFA1pr-*<br>*his3Δ::HPHX6 nup82Δ::NUP82 (S.*<br>*mikatae)-KANMX6* | This study |
| YPAR0147 | *MATa ura3Δ0 leu2Δ0 his3Δ1*<br>*can1Δ::LEU2+-MFA1pr-*<br>*his3Δ::HPHX6 nup82Δ::NUP82 (S.*<br>*kudriavzevii)-KANMX6* | This study |
| YPAR0149 | *MATa ura3Δ0 leu2Δ0 his3Δ1*<br>*can1Δ::LEU2+-MFA1pr-* | This study |
|  | *his3Δ::HPHX6 nup82Δ::NUP82 (S. bayanus)-KANMX6* |  |

